# Tracking and optimizing human performance using deep reinforcement learning in closed-loop behavioral- and neuro- feedback: a proof of concept

**DOI:** 10.1101/225995

**Authors:** Martin Dinov, Robert Leech

## Abstract

Reinforcement learning (RL) is a general-purpose powerful machine learning framework within which we can model various deterministic, non-deterministic and complex environments. We applied RL to the problem of tracking and improving human sustained attention during a simple sustained attention to response task (SART) in a proof of concept study with two subjects, using state-of-the-art deep neural network-based RL in the form of Deep Q Networks (DQNs). While others have used RL in EEG settings previously, none have applied it in a neurofeedback (NFB) setting, which seems a natural problem within Brain Computer Interfaces (BCIs) to tackle using end-to-end RL in the form of DQNs, due to both the problem’s non-stationarity and the ability of RL to learn in a continuous setting. Furthermore, while many have explored phasic alerting previously, learning optimal alerting in a personalized way in real time is a less explored field, which we believe RL to be an most suitable solution for. First, we used empirically-derived simulated data of EEG and reaction times and subsequent parameter/algorithmic exploration within this simulated model to pick parameters for the DQN that are more likely to be optimal for the experimental setup and to explore the behavior of DQNs in this task setting. We then applied the method on two subjects and show that we get different but plausible results for both subjects, suggesting something about the behavior of DQNs in this setting. For this experimental part, we used parameters suggested to us by the simulation results. This RL-based behavioral- and neuro-feedback BCI method we have developed here is input feature agnostic and allows for complex continuous actions to be learned in other more complex closed-loop behavioral or neuro-feedback approaches.

## 1. Introduction

### 1.1. Importance of sustained attention or vigilance

Vigilance is often defined as the ability to maintain alertness and attention over a prolonged period of time, possibly through boring, tiring and/or long tasks. For example, driving a car, flying a plane, being a radar operator, or a surgeon during longer procedures, or a security guard of a building or person, all require attention to be high so that false negatives do not occur (i.e. so that targets that should be detected or things that should happen do not happen) as well as to avoid false positives (mistakenly recognizing some bit of anatomy as tumorous, or mistakenly imagining or even hallucinating, due to fatigue, a bleep on a radar, or thinking a car is overtaking behind you when the car may be at a constant and safe distance). In either case, the subject involved makes a decision against reality, leading to at the very least a harmless realization of this, or some kind of minimal correction (e.g. realizing the car behind is not in fact overtaking), but it can also lead to catastrophic system failures leading to the loss of property, loss of life or financial losses, etc. A well-established fact of the ability of humans to sustain attention is that it is limited (Oken BS et al., 2006), and we are typically only capable of vigilance for short bursts of time typically, though we have some ability to prolong this period of sustained attention via top-down (more internally driven) strategies (Hopfinger, J.B. et al., 2000; Sarter, M. et al. 2001) and cues (e.g. reminding oneself throughout a task that it is important and you must refocus, recalling why it is important to you, or replaying certain mental strategies to keep distractions away, as well as noticing more or less consciously different timings and task-related conditions that can help in steering behavior, etc) but it is also possible to prolong sustained attention via bottom-up or more externally driven strategies or cues (e.g. playing a sound to remind of falling attention, or vibrating a car’s steering wheel, flashing a light briefly, etc). The problem with the top-down approach is that it is more likely to lead to fatigue and eventual abandonment of the internal strategy or cues people give themselves. Incorporating bottom-up externally cued alerts of various kinds can therefore present an approach to improving attention, especially when coupled with monitoring of attention via, for example, electroencephalography (EEG). While attentional control, whether driven through bottom-up or top-down mechanisms, will be affected by fatigue in any case, bottom-up approaches are, in some sense, more independent of higher-level motivation, interest or ability to control one’s attentional systems. Vigilance is one type of attention that has been heavily studied but there are other types of attentional systems that have been identified, which have relevance for sustained attention and vigilance-demanding tasks and which can be ‘hijacked’ and used to improve vigilance performance on tasks.

### 1.2. Arousal and orienting for vigilance improvement

The broad literature on attention has found that attention is best understood as a number of interacting, partially overlapping systems that work more or less in parallel to meet conflicting and varying task demands. Perhaps most relevant to NFB and the current work are overall cortical (and bodily) arousal as well as the orienting systems. Some researchers have referred to arousal as the global “energy level” of the brain (Portas, C.M. et al., 1998), referring to the idea that it reflects the global, or average, excitability of the cortex (or the brain generally). Other authors have suggested that arousal has local and more global aspects (Barry, R. J., 2007). Indeed, vigilance or sustained attention has often been referred to as ‘tonic arousal’ or ‘tonic alertness’ (tonic referring to longer-lasting, broader, more widespread) as opposed to arousal on a shorter time-scale which is more often called just arousal or ‘phasic arousal’ or ‘phasic alertness’ (i.e. arousal which is just a shorter phase) (Oken, B.S. et al., 2006). Arousal is in large part (but not entirely) dependent on norepinephrine (NE), released primarily from the locus coeruleus (LC) (Oken, B.S. et al., 2006). This LC-NE release affects the cortex and rest of the brain globally, as the NE neuromodulator is released to most brain regions in a non-specific manner (Petersen, S. E. and Posner, M. I., 2012). This NE release from the LC is likely to be largely controlled by non-cortical inputs from brainstem areas (in particular other neuromodulatory nuclei), hypothalamic areas, amygdala and some (perhaps lesser) contributions from the spinal cord and periphery via the vagus nerve, though there are also some important prefrontal cortex contributions (Oken, B.S. et al., 2006; Petersen, S. E. and Posner, M. I., 2012). Orienting can refer to either spatial orienting or temporal orienting, most broadly (Petersen, S. E. and Posner, M. I., 2012; Weinbach, N. & Henik, A., 2012). In either case, orienting is the act of the brain re-prioritizing current processing and (at least briefly) focusing energetic and attentional resources to a sudden new stimulus location/timing/feature set. In effect, this means that one area of the cortex or brain is given more blood, glucose, or whatever other resources are necessary to perform its temporarily increased need/recruitment compared to other areas of the cortex or brain. This is clearly important from an evolutionary perspective, as an animal (such as a human) must be able to prioritize mental and physical resources to a specific location or event if a certain external event/cue signals the animal/human to do so (e.g. roaring of a tiger, or a car in the distance alerting a human jaywalking despite a likely impending collision). The temporal orienting is when a cue (typically, and usually by definition, external) signals that an event is likely to happen in a specific (usually upcoming) time interval (Weinbach, N. & Henik, A., 2012). Spatial orienting happens in the brain when a (usually external) cue tells us that an event is likely to happen in a specific spatial location (Weinbach, N. & Henik, A., 2012). It seems that the temporal orienting system is more left hemispheric and the spatial orienting system more right hemispheric, with both orienting systems generally being most dependent/localized in superior parietal areas, close to areas associated with sustained attention, which likewise depends on parietal cortical regions. Both of the orienting systems seem to interact with the NE arousal system, and so they present us one way of modulating arousal in a bottom-up (and potentially completely automatable and consciously-effortless manner). This is important for NFB as we can therefore in part ‘hijack’ the brain’s natural arousal systems (whether temporal or spatial) with externally driven alerts or cues – and by monitoring and timing these cues to when the arousal/attention drops below a certain threshold (as estimated by certain EEG-derived measures for example), we can, to a non-trivial degree, maintain a certain level of arousal and vigilance.

### 1.3. Overview of RL and its use in this work

The term and field of reinforcement learning (RL), as used here, is a branch of machine learning (ML). It is similar to, but distinct from, supervised learning, which is perhaps the most commonly applied form of ML. In supervised learning, the correct target value at any given training round is always known and given to the learning algorithm explicitly. This is typically contrasted to unsupervised learning, where no target output values are given for the input values during training – in unsupervised learning the algorithm finds clusters and other types of patterns in the data directly. RL is only vaguely between the two – it is more like the former (i.e. supervised learning), with some features of the latter (i.e. unsupervised), but distinct from both with a rather different mathematical foundation on which it is based (Sutton, R.S. & Barto, A.G., 1998). We often refer to an RL system or algorithm as an “agent” that acts on the world (physical or virtual) in some way through actions. In RL, the main feedback that we provide to the agent is how well it performed, but not what the correct action or decision should have been in any given situation (unlike in supervised learning, where we will adjust algorithmic parameters precisely according to how far we are from the exact target value we were striving to output with the algorithm for the corresponding input for a given input-output pair). This leads to often more unpredictable outcomes – while the agent may learn to reach the general feature or goal, it may do so in highly peculiar, unpredictable and even undesirable ways, as we describe in the discussion section. However, this approach has recently proven very powerful in creating super-human game-playing capabilities, perhaps most notably in combination with neural networks by Google DeepMind (Silver, D. et al., 2016), though there are many other successful and inspiring examples of RL applications. In order for an RL agent to learn the desirability of a sequence of actions, we need to decide on the state-space that the agent can be in. Another way of understanding this is that we need to define the world in which the agent will see the states it is in and act while in those states, in order to reach other (more desirable) states. This can be done very explicitly or discretely, as with a lookup table/multidimensional array where each bin may refer to a specific state-action combination, or less explicitly and even continuously via function approximation (using e.g. linear functions or MLPs) (Sutton, R.S. & Barto, A.G., 1998). We use a full end-to-end Deep Q Network (DQN) as originally proposed in (Mnih, V. et al., 2014).

While there are other approaches we could have chosen for personalizing neurofeedback (NFB) and learning attention-related parameters for NFB, RL is quite fast/efficient in many cases, intuitively easy to understand conceptually and extremely flexible and general in its applicability – as such it is an ideal candidate for real-time and personalized learning in a complex stochastic environment requiring control actions to optimize some parameter(s) of interest.

### 1.4. Use of RL in previous EEG research

RL has been applied to EEG-based work previously with some success in a number of ways. In an early work in a somewhat relevant direction, (DiGiovanna, J. et al., 2009) suggested and implemented an adaptive (what they called a ‘co-adaptive’) RL-based BMI system where the system learns from the user’s (in their case a rat’s) motor cortex signals and the rat adapts to the system’s outputs and the goals via implicit reward/punishment signals. No less relevant to this work is (Iturrate, I. et al., 2010), where the authors used specific aspects of the user’s own EEG as a reward signal and learning is done using these EEG-derived reward signals in a virtual BMI setting with two human participants. The rewards were derived by classifying parts of the EEG outputs into classes, which were then used as the feedback/learning reward signals. Interestingly, they show some amount of generalization across (similar) tasks with the RL system. (Pohlmeyer, E. et al., 2014) developed an actor-critic-based RL system for brain machine interface (BMI), using marmoset monkeys and intracranial microelectrode array in the motor cortex to control a robotic arm, being given rewards in the form of food (and therefore not explicitly encoded and precisely controlled reward amounts). Another good application of RL in a relevant but quite different domain is (Moore, B. et al., 2014) who used RL for creating a closed-loop anesthesia control system. While the domain of their work is not within BCI or BMI, they make a number of good points and observations, discussing in some detail their RL implementation details, however their system was not *adaptive,* as it did not learn during control (where control in their case would probably have been varying the amount of propofol given to subjects that were being anesthetized). They discuss the reasons for this, citing that the lack of a predictable and unchanging policy, while more flexible and subject and situation-specific makes it harder to get approval for use of such adaptive control systems in a clinical setting. As the current study is more a proof of concept and we wanted to explore various aspects of EEG and attentional control using RL in some detail, we opted for an adaptive control-based learning. Another relevant work was [4], who used a variant of RL for improving clustering on a motor imagery data set, but not in a real-time or control/learning setting. More recently (Wang Y. et al, 2013) used another form of RL (AGREL) for modeling and predicting in a BMI setting with a primate, though their RL system was not used for learning from the brain signals, but rather from the motor effects/movements generated. To the best of our knowledge, RL has not yet been applied in a purely neurofeedback context, which is what we have succeeded in doing and present here. While the system we implemented has a number of easily generalizable features that overlap in large part with previous work, we also explored in some further detail the parameter-dependence on the system in a simulation setting, unlike most of the previous works cited in this section. We also suggest that NFB, in particular attention monitoring and control, is a very natural and efficient setting in which to apply RL in the future – e.g. for the portable (and personalized) monitoring and improvement of attention via feedback systems (e.g. in cars, planes and so on). Finally, we are the first to apply state-of-the-art Deep Q Networks (DQNs) for the reinforcement learning in this context. DQNs have proven very powerful in other domains, where super-human performance has been achieved using self-learning via reward signals. We adapted a Sustained Attention To Response (SART) implementation to be compatible with the OpenAI gym Environment API (Brockman G. et al., 2016), which allowed us to use a gym-compatible RL framework built on top of the Keras ML framework called Keras-RL, which includes various types of DQN agents.

### 1.5. Previous work in personalization and real-time learning for neurofeedback

There has been some amount of work done with neurofeedback, using various modalities. With FMRI, some recent real-time and personalized neurofeedback and BCI work includes (Sherwood M.S. et al., 2016), where real-time personalized feedback was successfully demonstrated across different scanners strengths and across multiple task types with 10 subjects. (Lorenz R. et al., 2016) developed a novel type of neurofeedback technique using Bayesian optimization to automatically tweak and generate task parameters to get a desired activation in given brain regions (i.e. a desired brain state) in a proof of concept study. In a simultaneous EEG-FMRI setting, (Zotev V., et al., 2016) used EEG to monitor electrical activity changes during a realtime FMRI neurofeedback study of the amygdala. They found frontal EEG asymmetry changes to correlate to amygdala regulation-related BOLD changes, suggesting that the EEG signature could potentially be used as a proxy measure of amygdala activity regulation. More directly relevant to the present work however is (Zich C. et al., 2016), who did EEG-based real-time NFB in a simultaneous EEG-FMRI setting in a motor imagery (MI) setting. Interestingly, they showed that while contralateral FMRI activations in sensorimotor areas did correlate to contralateral EEG activation changes during the real-time EEG NFB in the MI setting, ipsilateral BOLD activity changes during the NFB in sensorimotor areas did not correlate to ipsilateral EEG changes. They used signals at high GFP points and applied Common Spatial Pattern filter followed by LDA classification for left and right-hand MI. (Iacoviello D. et al., 2015) used EEG in a real-time BCI classification study of self-induced emotions and seeing if a single channel can achieve similar accuracy in identifying one particular emotion (disgust) compared to using all channels. They used a number of methods but ultimately classified using a Support Vector Machine (SVM), showing that the single channel changes can identify at least that one emotion. An adaptive approach was taken in (Hsu S. H. et al., 2016), who developed a optimized online recursive ICA algorithm (ORICA) using online recursive least squares (RLS) for real-time blind source separation (BSS), which is of obvious interest and use to BCI. Another interesting study was (Luu T. P. et al., 2016), who used closed-loop EEG-based BCI system for controlling the gait of a virtual avatar in VR in a BCI-VR setting. They used slow cortical potentials (0.1-3Hz) activity, achieving R values of around 0.4 for joint control of the virtual avatar on average. These numbers are typical accuracies observed in non-trivial BCI control settings.

The cited studies are a small subset of the types of BCI and neurofeedback that have been done. For NFB specifically, where the system learns in real time from the individual (and is thus personalized) depends on the user seeing their current brain activity in some way. This, while useful and successful, requires significant effort by the user and distracts from being able to do an actual task in parallel. One can also do NFB without requiring the user to do anything by passively having them perceive certain stimuli (typically sound or light cues). This latter approach can be referred to as passive NFB versus the former (where the user has to exert effort), which we can call active NFB. Our work presented here provides a very general purpose and powerful approach to performing passive NFB, though the approach need not be restricted to passive NFB and is probably useable in all BCI settings.

## 2. Methods

### 2.1. Reinforcement Learning framework

We wanted to apply end-to-end RL that is close to state-of-the-art. We used the Python-based Keras-RL framework which runs on top of Keras. Keras-RL implements a number of deep reinforcement learning methods in a flexible way using Keras, allowing for easy configuration of the number of, sizes of and types of the layers in the underlying neural net. We used a Deep Q Network, or DQN – this type of deep end-to-end RL allows for a full modeling of the state and action spaces, directly modeling the desired behavior and learning of the agent with reference to the modeled environmental dynamics – i.e. there is a single deep (multi-layered) neural network that is trained fully by the reinforcement signals continually received by the environment, which are a function of the subject’s EEG and behavioral (reaction time) output.

In our case, the environmental dynamics were EEG signals acquired by a portable, low-cost commercial EEG device (see section on EEG acquisition below), as well as the reaction time (RT) as the behavioral signal fed into the RL agent. The RT is also the basis of the rewards given to the agent. We tried a number of learning rules, with the two most successful ones both being based on a moving window averaging of the recent RT of the subject doing the task (see below for the task details). We waited for n number of trials before we had enough samples to begin computing the mean RT from the recorded behavioral responses, giving rewards of 0 until that point. We chose n=10, but also tried n=3. While n=10 takes slightly longer to start learning, we find that n=3 did not work as well in practice or in simulation (see corresponding sections for details). So, with n=10, on the 11th point we check whether the current (11th) RT is greater or smaller than the mean of the last 10 RTs. In the first version of the two main rewarding mechanisms we investigated, we rewarded according to the difference between the current RT and the last 10 RTs, which are computed in a moving window fashion as new RTs come in during the task, i.e.:

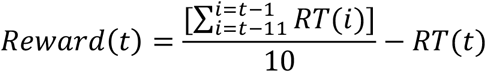

In the second version, the reward is a 1 if the current RT is lower than the previous 10 RTs and -5 if it is greater than the previous 10 RTs:

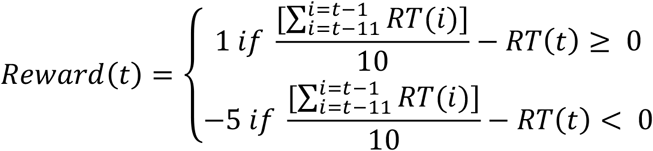

We originally used identical rewards/punishments (0.3 and -0.3), closer to the actual RTs, as this should lead to a faster convergence, but these led to the agent getting stuck in one of the two actions for long intervals of trials.

In order for the agent to explore the state-action space sufficiently well and find ‘good’ solutions, the agent needs to not get stuck in any one action for too long. There are different exploration strategies that can be used. We explored the use of *ε*-exploration and Boltzmann-exploration for both in simulation and the experiments ran. In the former, we select the best (so far found) action (i.e. alerting or not alerting) with probability 1-*ε*. We select one of the two actions uniformly with probability *ε*. This means that the agent is acting greedy (selecting the best action found so far) with an overall probability of 1-(*ε*/2). As long as *ε* is not too small, the other action will be chosen often enough for the agent to realize that it is (currently) a better one to choose. In Boltzmann exploration, we have a softmax or Boltzmann distribution-weighted selection of probabilities for selecting a given action *a* from all possible actions:

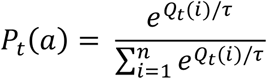

In our case, we have:

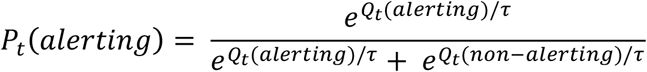

The *τ* parameter is a temperature parameter that is typically annealed over time and controls the degree to which the actions are weighted according to their Q values. A high temperature parameter leads to more random action selection and thus more exploration, while a small *τ* leads to behavior closer to greedy action selection (we have fully greedy action selection for *τ* → 0). In between, the agent explores non-optimal actions with some probability that depends on the specific range of the Q values. Since this depends on the rewards used, *τ*/Boltzmann-action selection is more context-sensitive than *ε*-exploration. In general, we can imagine that Boltzmann exploration leads to better outcomes as the non-optimal/non-greedy actions are weighted according to their estimated Q-values (i.e. according to how good the agent thinks those actions are), which is a more informative action selection than uniformly randomly selecting among all other actions with some probability ≈ *ε*. In practice, finding a good *τ* is more difficult and less intuitive than finding a good *ε*. We explored the use of both on this type of BCI/NFB presented here.

There were only two actions that we used, for simplicity of analysis. Action 0 is no alerting, i.e. doing nothing in that trial and action 1 is alerting using an auditory phasic alert. While we could also train on other parameters such as the strength/volume of the alerting cue, or the phase in the specific band of the EEG, these make interpretation of the agent’s decisions and the subsequent results even more complex and do not add to the proof of concept work presented here.

### 2.2. Task and cognitive setting

We used the Sustained Attention to Response Task (SART), which has previously been used in a number of seminal studies of attention. It requires the subject to press a given key (we set this to the space key) on all numbers seen except for a specific given number (we set this to be 3). This task requires both alertness, selective attention and inhibition control, as well as sustained attention when done for sufficiently long. After each response, very briefly a neutral visual stimulus is presented (i.e. a mask). The requirement of the user to withhold keypresses on visual stimuli (the number 3) means that the task is not easily automatable without incurring significant errors. By penalizing any errors significantly in the RL agent, we take into account the error rate during the task. Either a false positive (withholding a response when it shouldn’t have been withheld – i.e. not pressing a key when the stimulus was not a ‘3’) or false negative response (responding when a response should have been withheld – i.e. pressing ‘b’ when there was a ‘3’ and no keys should have been pressed) was penalized as a high reaction time of 1 seconds (1000 milliseconds) – this roughly corresponds to the slowest reaction times seen by the two subjects on any trial, during training on this implementation of the SART and during the actual experimental runs that are presented and discussed here. Since the mean RT was around 250-300 milliseconds both in experimental data and in simulation data, even with the heavy right tailed distribution of RTs observed experimentally and then modeled in the simulated data, 1 second constitutes a ‘very bad’ response, pushing the mean of the moving window considerably higher. We chose to do this to take into count the error rate, which is known to increase in more alerted states. The purpose of this kind of behavioral neurofeedback is to ultimately improve performance on a given task and decrease error rate as well. In other words, while alerting leads to lower reaction times in general, we also wanted to improve on accuracy/error rate. While shorter RTs resulting from the alerted state are an ‘improvement’, a significantly higher error rate may not be acceptable, depending on the real-world task. We can think of the high reaction time penalty as a kind of regularization applied to the loss function of the RL system.

The experiment was programmed in Python using the open source library Expyriment (Krause, F. & Lindemann, O., 2014).

### 2.3. Portable EEG acquisition

In order to simulate sensible EEG and behavioral (reaction time) data, we could have referred to the literature and simply pulled out standard values. Indeed, this seems like a reasonable approach and we did that too, but we also ran the SART on two healthy adult subjects, while recording their EEG frontally from Fp1 and Fp2 using the InteraXon Muse headset. We chose this commercial EEG headset as it is readily-available, affordable and easy-to-use. It is one of the most widespread consumer EEG devices and we are interested in creating a neurofeedback BCI system based on such widely-available devices, so as to increase the usefulness of the developed methodology. While laboratory results using high-density EEGs provide valuable scalp coverage of the brain’s electrical activity, many phenomena, including attention, have EEG components that can be picked up from specific scalp locations – such particular use cases do not require fuller scalp EEG coverage. The headset has different sampling and signal processing capabilities built into the hardware and associated software. We used the Muse-IO application provided by InteraXon, sampling the electrodes at 3520Hz and sending data at a downsampled 220Hz, with a notch filter set to 50Hz to remove line noise. The Muse uses a Driven-Right-Leg circuit for the reference and ground electrodes which is used to reduce common-mode noise from the recording electrodes (Fp1 and Fp2). Finally, we derive relative band power values computed by the Muse-IO application in the delta (1-4Hz), theta (4-8Hz), alpha (7.5-13Hz), beta (13-30Hz) and gamma (30-44Hz) bands. The ranges are inclusive and overlapping frequency bins are counted towards both neighboring bands (e.g. the last bin at 4Hz is part of both delta and theta). The relative band power values are computed from the log-scale band power, which in turn is the sum of the power spectral density (PSD) of the EEG data in the respective frequency interval.

The linear band power values are computed as:

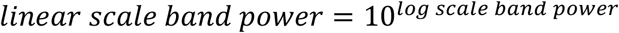

Using the above, for the relative alpha band power (proportion of the PSD-computed band power that falls in the alpha band range) we have:

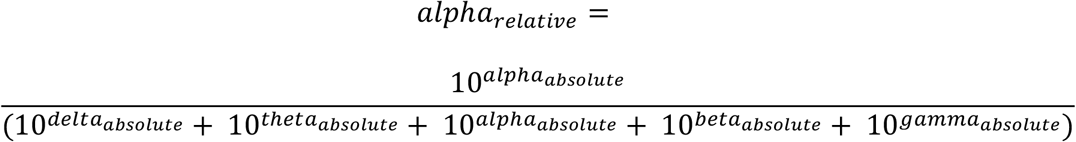

The other relative band powers are computed analogously, as the relative proportion from all the bands of the linear scale band power. This gives us values in the range [0,1] which are computed in a sliding window fashion and emitted by the Muse-IO tool at 10Hz.

### 2.4. Simulation

The process of simulating the data that we used was driven empirically. We used one of the two subjects to do a run of SART with 300 trials, with 33% of them randomly alerting, to get some indication for the difference of reaction times for this subject between alerted and non-alerted states, as well as the distribution of the EEG band powers. Figure 1 shows the recorded EEG band values, the reaction times and the corresponding fitted lognormal distributions. Because the EEG bands did not display obvious differences between alerted and non-alerted states in themselves, we show only the fitted distributions and plots of both alerted and non-alerted states combined, and not individually. Note also in Figure 1 that a lognormal fit seems to be a decent fit to the empirically-observed reaction times from this subject. We tried a number of distributions and found that a lognormal distribution is the single distribution that tends to best model the different bands as well as the reaction time. We discuss some issues and implications of this choice. However, the variability of the band powers and the reaction times are such that slight deviation from the distributional form chosen is not going to significantly alter the shape of the distribution, whether a lognormal fit is ‘good’ or not. As such, the lognormal fits were deemed sufficiently good for generating simulated values with the corresponding distribution parameters found from subject 1’s non-feedback SART data. These are: *μ* = −1.23 and *σ* = 0. 293 for the relative delta power lognormal fit, *μ* = −1.65 and *σ* = 0.295 for the theta, *μ* = −1.73 and *σ* = 0.368 for the alpha, *μ* = −1.69 and *σ* = 0.258 for the beta and *μ* = −2.22 and *σ* = 0.318 for the relative gamma band power distribution. We find maximum likelihood estimates of lognormal distribution parameters of *μ* = −1.09 and *σ =* 0.175 for the alerted reaction times and *μ* = −1.17 and *σ* = 0.175 for the non-alerted reaction times.

**Figure 1:**
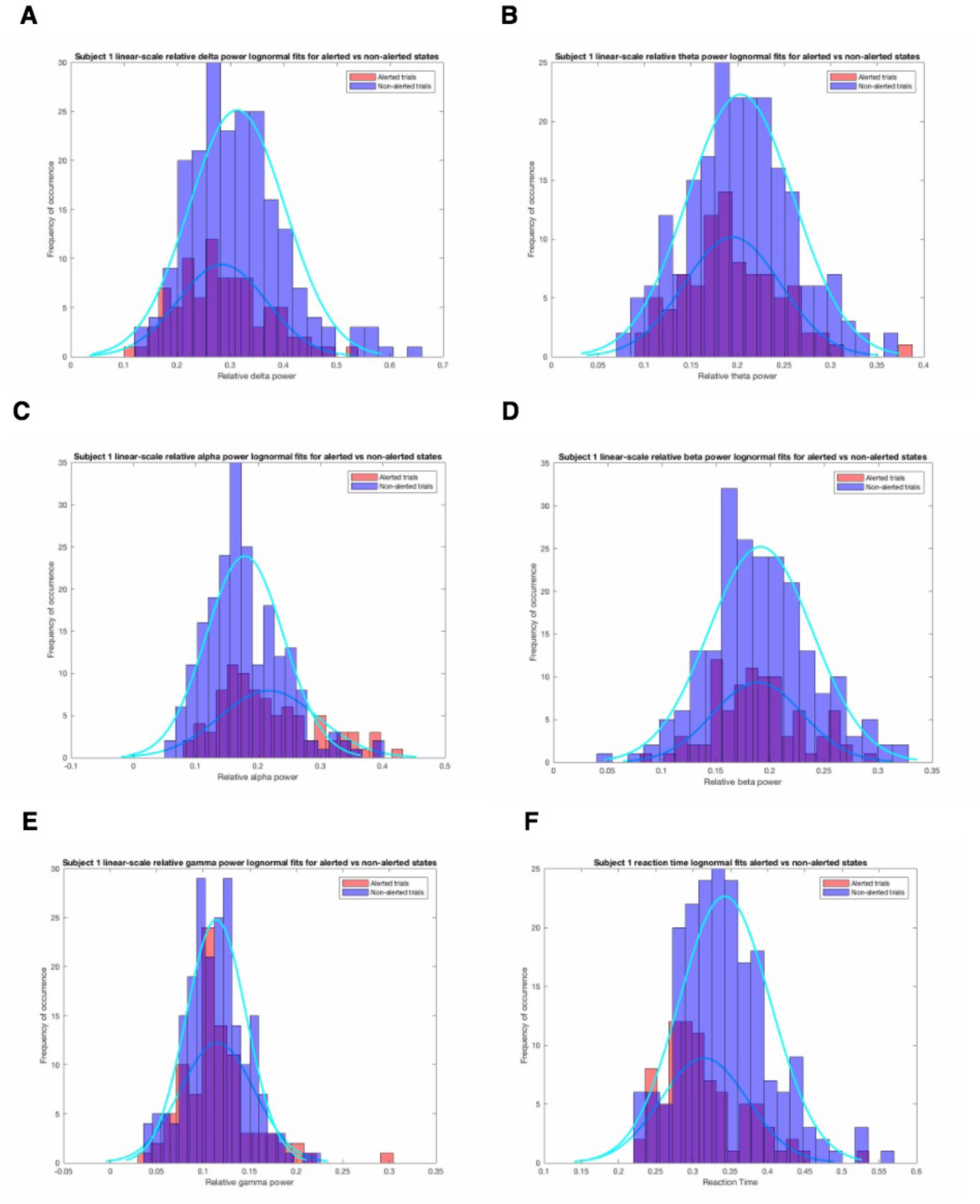
Histogram plots and lognormal fits based on the band power data from subject 1 performing SART with 300 trials and randomized alerting in 33% of the trials (sub-figures A-E) and a histogram plot and lognormal fit of the corresponding reaction times from subject 1 for alerted (in red) and non-alerted (in blue) trials. Note that for the reaction times, we excluded 19 trials which were incorrect, as we penalize and model an incorrect response as having a reaction time of 1 second, which would unnecessarily skew the corresponding fitted distribution of the reaction time to the right. While the incorrect trials tend to be very fast responses where response inhibition did not suppress the incorrect response, such incorrect responses can be costly in various real-world situations and the way they are modeled can strongly influence the real-time adaptive learning system.

Using the MLE parameter estimates of the reaction times, we then simulated values from those distributions to feed more empirically-tuned values for this task and this subject to the DQN simulation, in order to tune the parameters of the RL learning to where it learns successfully and quickly. Figure 2 shows the simulated distributions using the subject 1’s empirical data.

**Figure 2:**
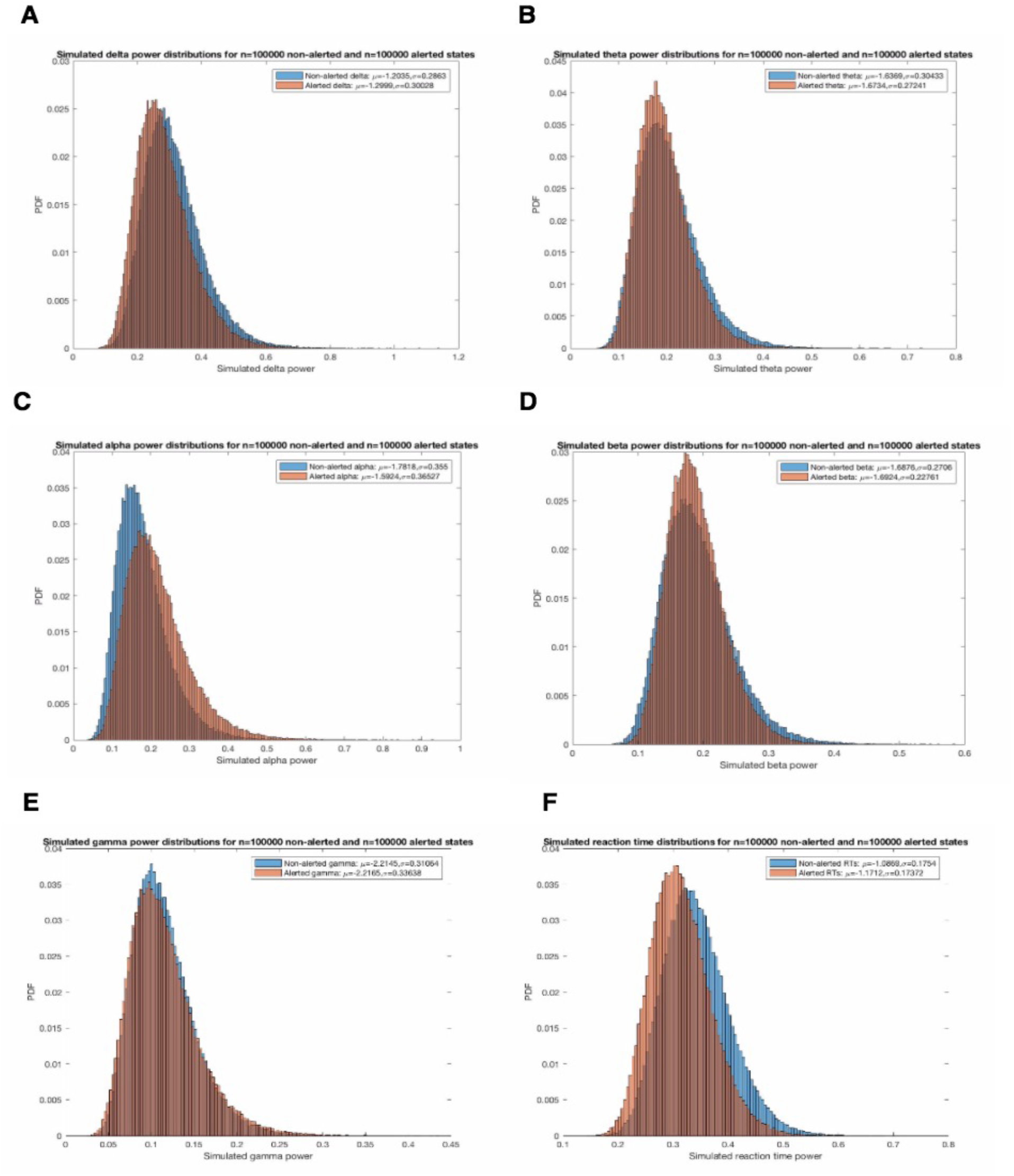
The figure shows lognormal simulated data with parameters (*μ* and *σ*) derived from subject 1’s 300-trial SART results shown in Figure 1 (without behavioral/neuro-feedback with random alerting). Most noteworthy is that relative frontal alpha power is significantly higher in alerted states than in non-alerted states (Figure 2C). Most other relative band power measures do not differ as noticeably, though relative theta power is lower in alerted than non-alerted states. The simulated reaction time values, driven by the actual reaction time values from the subject, show that alerted states, while having considerable overlap with non-alerted states, tend to have lower reaction times.

### 2.5. Experimental setup

The two subjects then performed 5 runs of the SART each. Both subjects had previous practice on this implementation of SART to familiarize themselves with it. The task was performed at night time when the subjects were already somewhat drowsy, to increase the need to pay attention to the task and to hopefully increase the contrast between non-alerted and alerted states.

The alerting cue was a 3000Hz sound generated as a 3000Hz sine wave in Audacity. The sound file was saved as a pulse coded modulated uncompressed wave file sampled at 44.1 KHz with 16 bits per sample, played at maximum volume on the same computer system, with the same set of Audio Technica ATH-M50x headphones plugged into the audio/headphone jack. The sound was loud enough to initially be described by both subjects as startlingly loud.

### 2.6. Analysis

We used the Student’s 2-sample t-test on the reaction time results for the final results. We did also use the Wilcoxon signed-rank test as a non-parametric version of the t-test, as well as the unequal-variance t-test (Welch’s t-test), though they gave almost identical results in all cases, so are not reported in the results. Finally, we ran general linear models (GLMs) for each condition, testing whether the RTs were at all predictable by the relative EEG band powers, using linear terms, but also including interactions between the features/regressors.

## 3. Results

### 3.1. Simulation Results

We first report here on a sub-set of the results of running simulations, showing some of the effects of the most important parameters in the system. In Figure 3 below, we show some of the effects of tuning two of the DQN parameters (*ε* and *γ*) in the learning process, using *ε*-greedy exploration of the action space with 3 ReLU hidden layers, each of width 4 neurons, with a final linear activation layer with 2 neurons for the two actions (alerting vs non-alerting).

**Figure 3:**
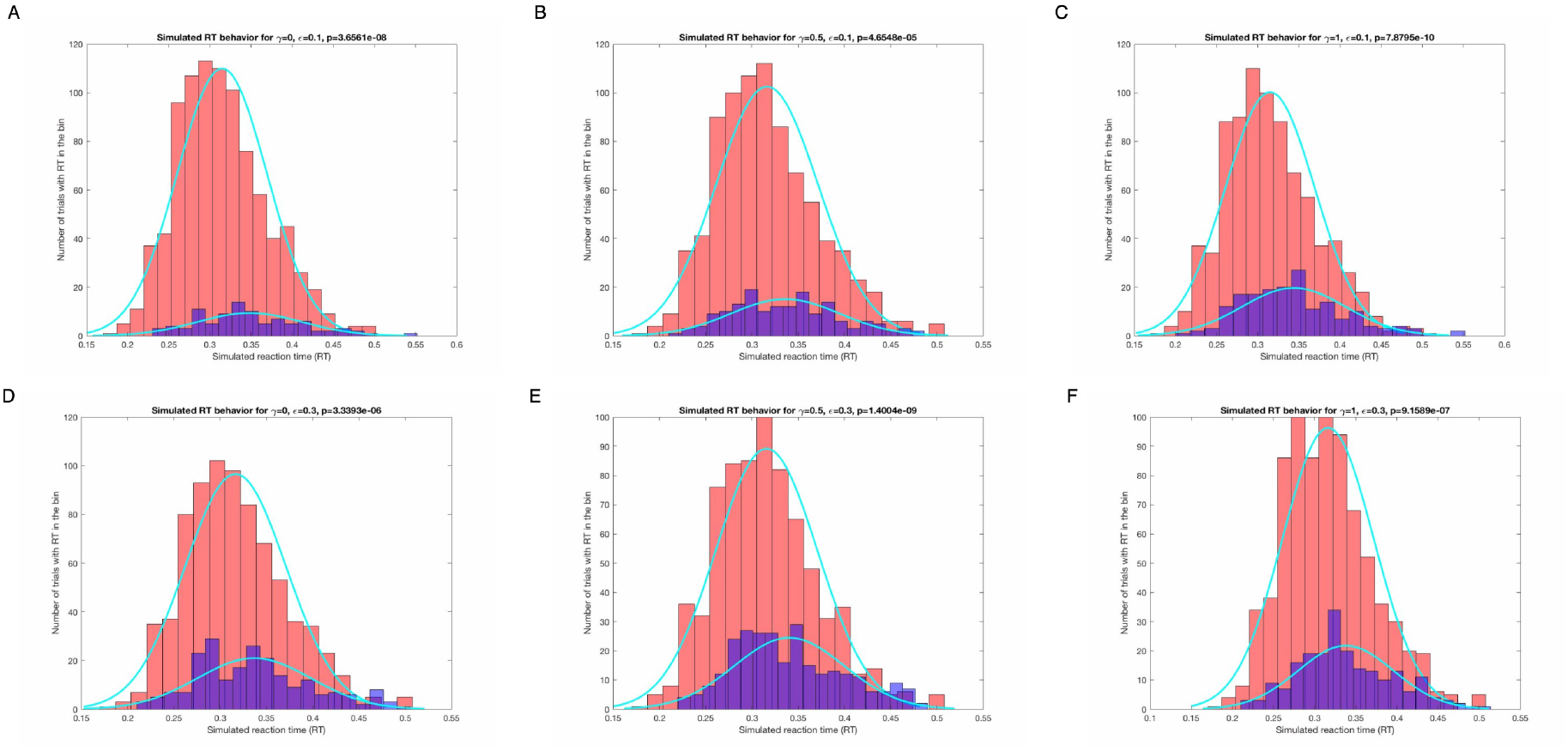
The figure above shows the learned results of the difference in the two distributions (alerted, in red, and non-alerted trials, in blue) for differing values of *ε* and *γ.* We also computed 2-sample t-tests in each case, showing the p-values in the title of the respective sub-figures. The t-test p-values give an indication of the degree to which the agent was able to learn the two different distributions.

Note that in Figure 3, as expected, increasing *ε* leads to a higher proportion of nonalerted trials, since more trials are picked uniformly randomly (with *ε =* 0.3 in the bottom three figures). While we show the case of *γ* = 1 in the above figure, we do not show this in Figure 4 below or explore the use of *γ* = 1 in the experimental results, as weighting all rewards from indefinitely far in the future (or past) does not make much sense, especially in a finite task setting. In Figure 4 below, we show results of gradually increasing the *τ* parameter in Boltzmann/softmax action exploration strategy. Note that we do not show the case for *γ* = 1 in Figure 4, but only show histogram plots of the achieved states with *γ* = 0 and *γ* = 0.5 for alerted (again in red, as in Figure 3) and non-alerted (again in blue, as in Figure 3):

**Figure 4:**
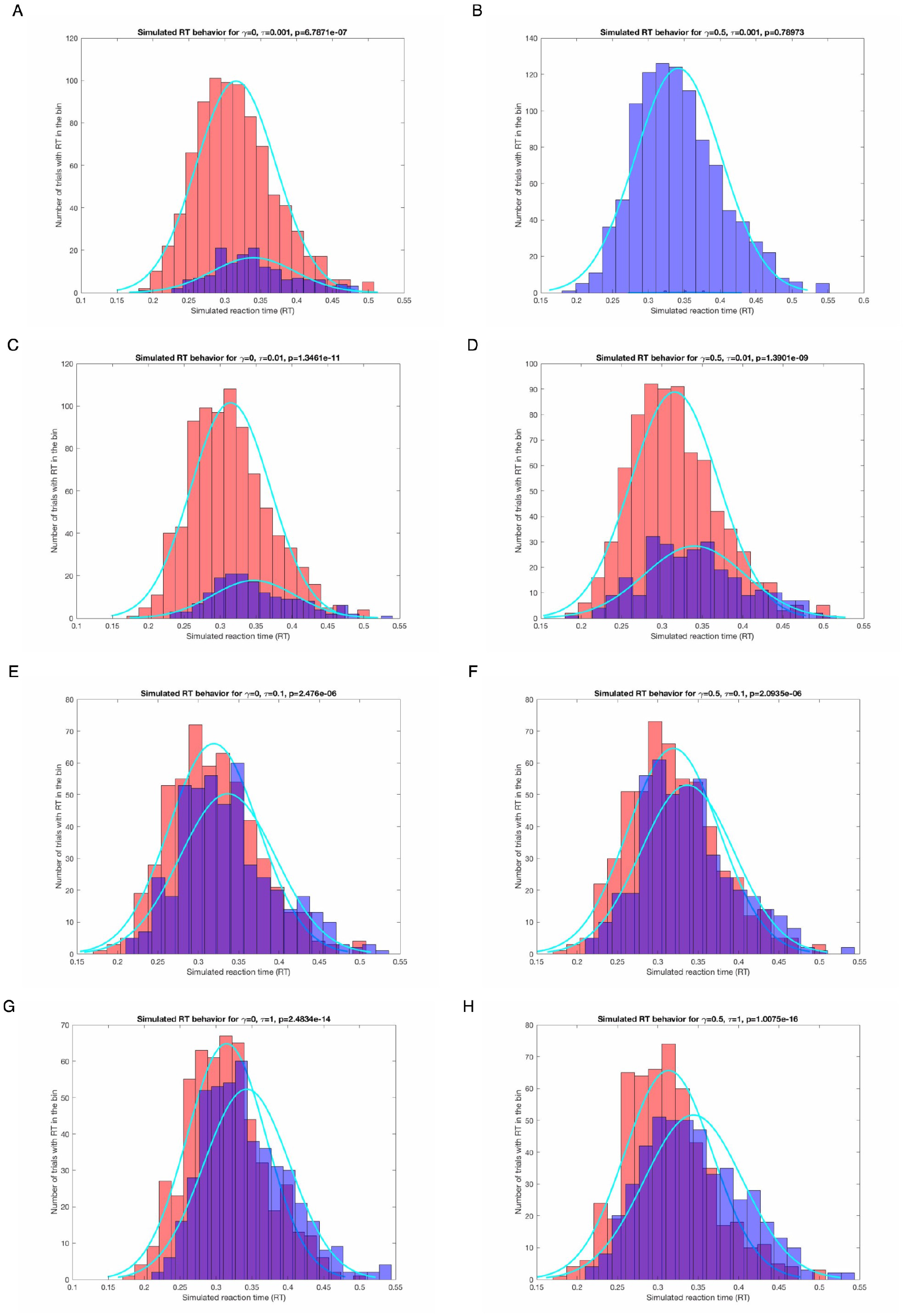
The sub-figures show histogram plots of the achieved states (reaction times) for values of *γ* = 0 and *γ* = 0.5 for four different sets of values of *τ* (0.001,0.01,0.1,1) in Boltzmann or softmax-driven action space exploration. Note that with this action exploration strategy, the agent finds that the alerting and non-alerting actions are much closer to being equally good than the *ε*-exploration strategy would suggest. Since with Boltzmann-exploration, given suitably chosen values of τ, we weight the different actions based on their utilities (i.e. their estimated Q values), the agent behaves less binarily in its choice of action. This is one of the main reasons we decided to use Boltzmann-exploration action-selection for the experimental runs.

As mentioned in the caption of Figure 4, Boltzmann-exploration leads to a more balanced choice of which action to take, due to its weighting of which action to select based on the estimated action utilities (i.e. the Q values). Figure 5 below shows four examples of the agent learning at different rates and learning different action preferences for different exploration strategy parameter choices:

**Figure 5:**
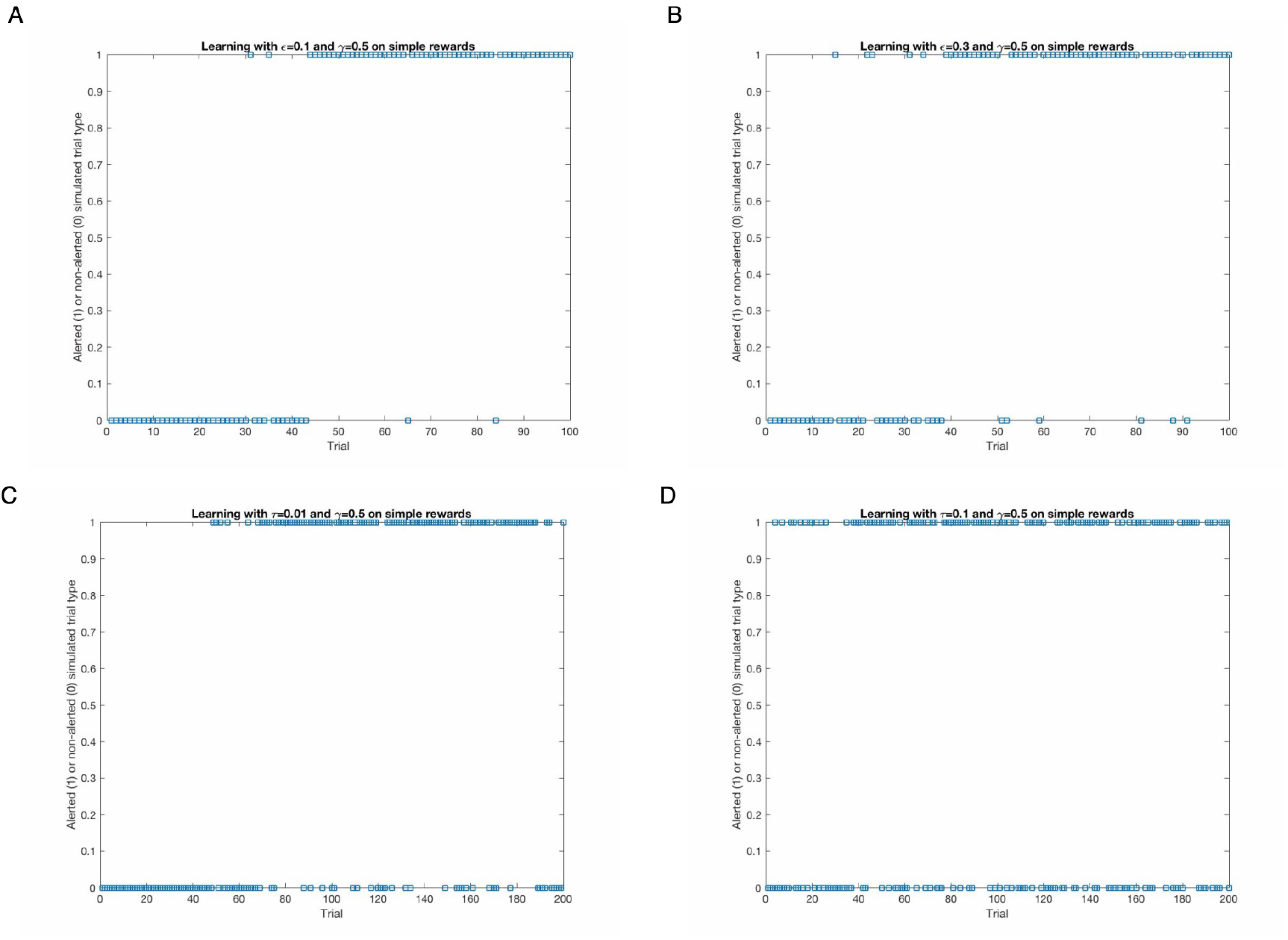
The top two figures (A and B) show the same agent (i.e. same architecture) learning the simulated reaction times for *ε* = 0.1 and *ε* = 0.3. Notice that while the agent in A (*ε* = 0.1) learns somewhat slower, it makes less non-alerting actions, as they lead to an overall higher reaction time. The agent in B learned a few trials faster to quasi-greedily prefer the simulated alerting state, but it makes more exploratory (non-alerting) actions. This is less useful in the case of stationary simulated data, but is important for real, and especially brain-driven, inputs. The bottom two figures show the learnings of Boltzmann-exploration-driven agents. The agent shown in C learned to prefer action 1, or alerting, after more trials than the agent in A, and continued to explore more after finding that it is the preferable state. The agent in D displays similar but even more exploratory behavior than C – still preferring alerting over non-alerting but even less so. Finally, note that all the simulated results shown here and discussed are of the same pseudo-randomly generated sequences which were always seeded with the same seed.

In simulation, such a graded action selection may or may not be useful, depending on the purposes of the simulation. Because both brain dynamics in general and EEG signals specifically are neither stationary nor as simple to model even on short time scales, we expected softer action selection using Boltzmann exploration of actions to prove more successful than an *ε*-greedy strategy.

### 3.2. Experimental Results

The figure below shows the RTs from all 5 experimental runs for subject 1:

Figure 7 below shows the RTs of subject 2 for the same conditions as subject 1:

**Figure 6:**
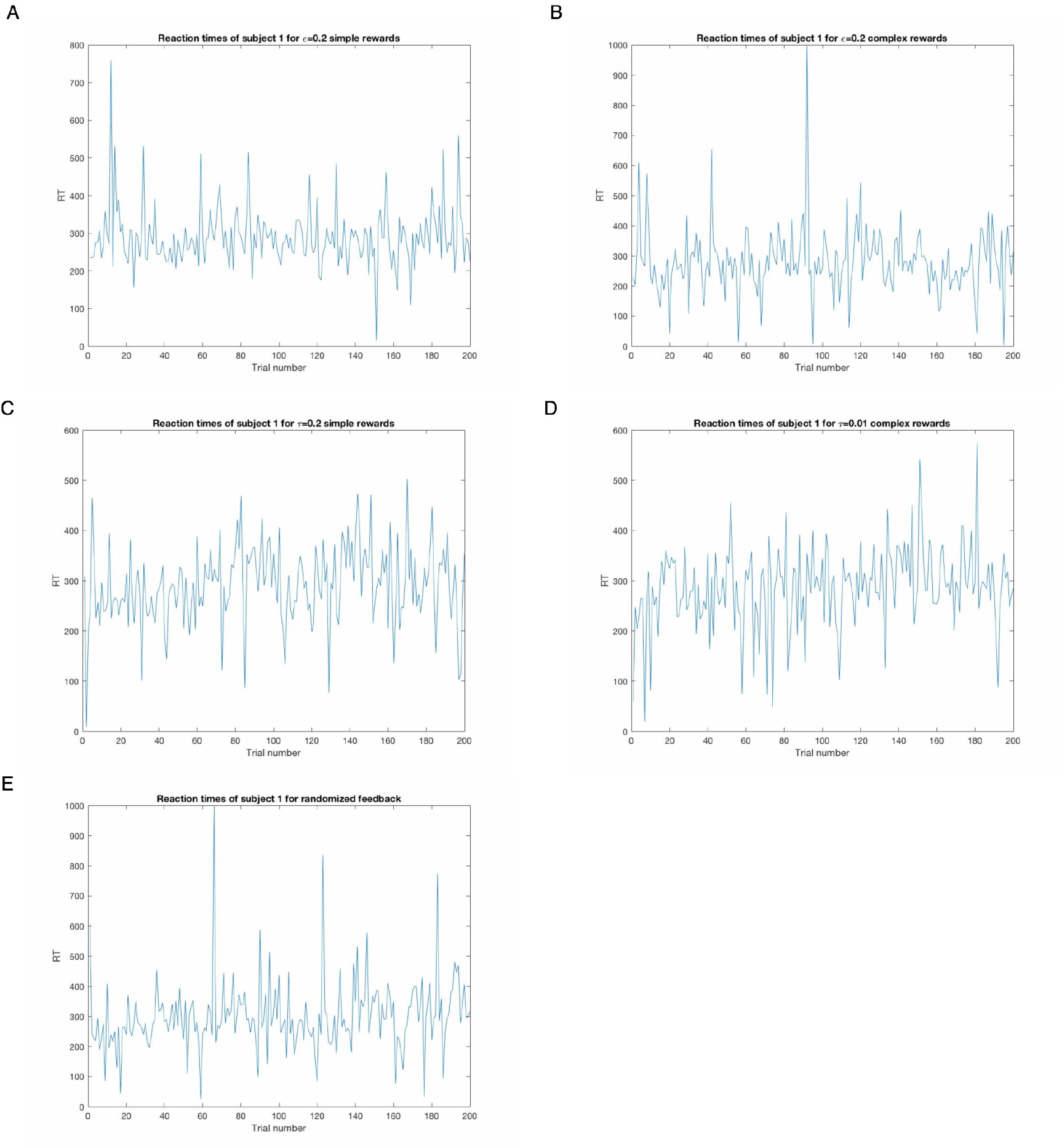
The figure shows the reaction times (RTs) of subject 1. Sub-figure A shows RTs for RL-driven NFB with *ε* = 0.2 with the simple reward rule. B shows the same *ε*-exploration with complex rewards. C shows simple rewards with *τ* = 0.01. D shows complex rewards for *τ* = 0.01. Sub-figure E shows RTs for the randomized alerting without RL-driven NFB (with probability of alerting = 0.2). Subject 1 missed responding to 1 trial in B and in 1 trial in E.

**Figure 7:**
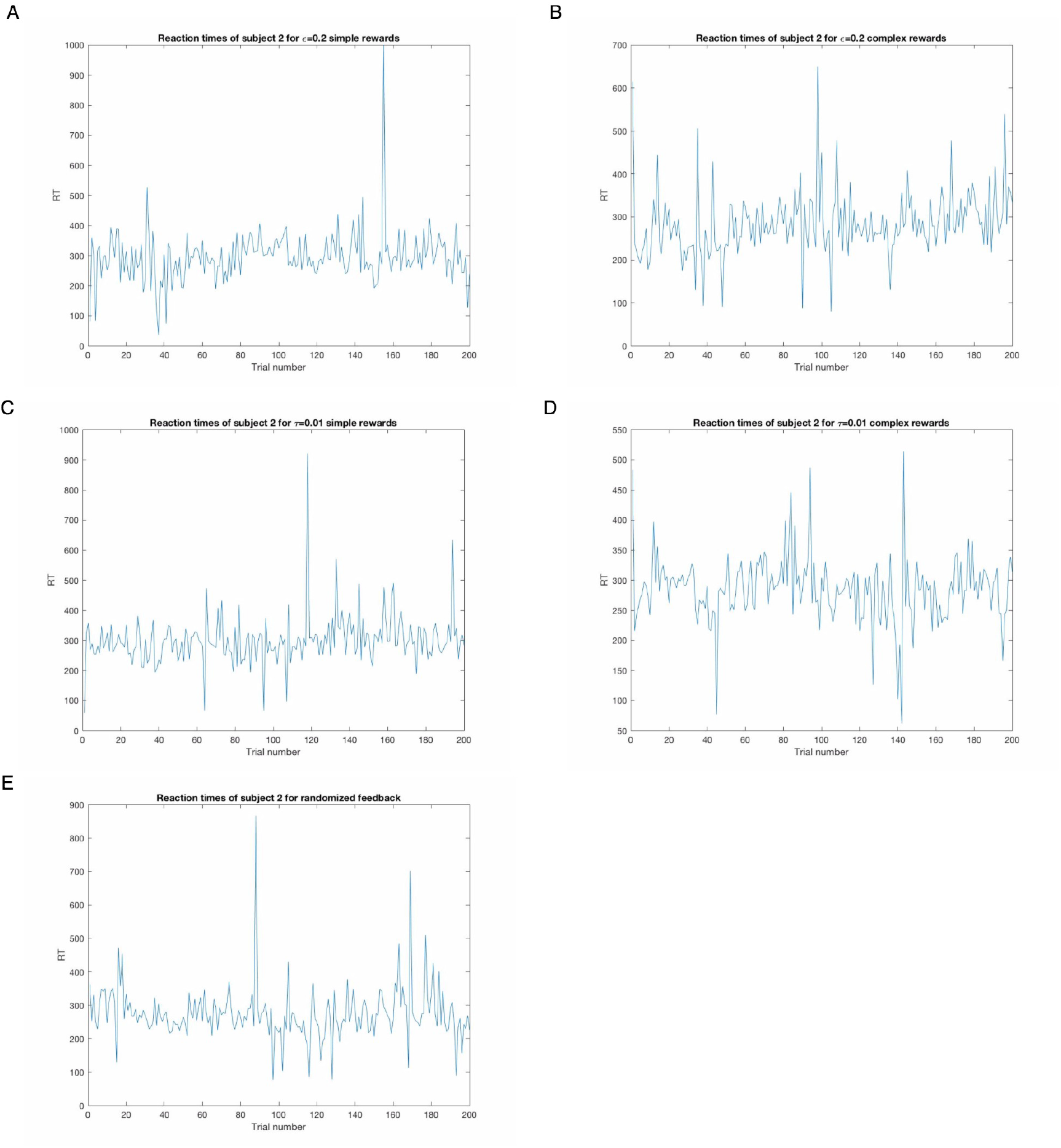
The figure shows the reaction times (RTs) of subject 2. Sub-figure A shows RTs for RL-driven NFB with *ε =* 0.2 with the simple reward rule. B shows the same *ε*-exploration with complex rewards. C shows simple rewards with *τ* = 0.01. D shows complex rewards for *τ* = 0.01. Sub-figure E shows RTs for the randomized alerting without RL-driven NFB (with probability of alerting = 0.2). Subject 2 failed to respond in 1 trial in A when they should have.

The reaction time plots themselves are not very informative in themselves – and ‘natural’ variability in task performance between tasks is going to have strong effects beyond that of the alerting frequency and timing. Based on just the reaction times, the simple rewards may have been more effective for subject 1, who had higher standard deviation of RTs during the complex rewards-driven alerting (across all reaction times, i. e. for both alerted and non-alerted trials). For subject 1, we found no significant difference between alerted and non-alerted trials for the *ε*-simple rules alerting (p=0.165). *ε*-complex rules alerting led to just barely significant (NB: without correcting for the multiple testing) results, with faster alerted trials than non-alerted (p=0.019). Subject 1 had very slightly faster RT on alerted trials than non-alerted trials (p=0.0302) for the *τ*-simple agent and a smaller (and ‘non significant’) difference for the *τ*-complex agent (p=0.114). Subject 1 had very similar RTs between alerted and non-alerted trials in the randomized (non-RT-driven) condition E (p=0.8834). Note that alerted states had lower RTs (and correspondingly lower p-values, sometimes significant when not corrected for multiple comparisons) for the RL-driven alerted states compared to the non-RL-driven randomly alerted states, shown in E. Subject 2 responded differently to the learned alerts for them. For the *ε*-simple case, subject 2 did not respond well to the alerts, having a somewhat higher RT on alerted trials than non-alerted trials (p=0.206). The *ε*-complex condition had almost statistically similar RTs (though note that there were only 35 non-alerted trials, p=0.3661). The *τ*-simple condition led to a small significant difference with faster alerted trials than non-alerted trials (p=0.0891). The *τ*-complex case for subject 2 led to no ‘significant’ difference, but there were only 7 non-alerted trials, so the t-test (and the other tests tried, but not reported) are not going to pick up the small difference in alerted vs non-alerted trials that we find in this task. Finally, the randomized (non-RL-driven) condition for subject 2, shown in E, led to strongly similar RTs between alerted and non-alerted trials (p=0.9302). Again, note that nonrandomized (RL-driven) alerting did lead to more significant differences between alerted and non-alerted trials.

Figure 10 above shows the weights from the input layer to the first hidden layer for all 5 conditions for subject 1. Figure 11 below shows these for subject 2. The columns are the different inputs (i.e. features) fed to the neural network underlying the DQNs: input 1 is the RT from the last trial and the next 5 inputs are the linear relative band power values over the last second for the delta, theta, alpha, beta and gamma bands, respectively. Sub-figure A shows these weights for *ε*-simple, B for *ε*-complex, C for *τ*-simple, D for *τ*-complex and E for the randomized (non-RL-driven) alerts that played about 20% of the time. As noted in the caption of Figures 10 and 11, the subfigures E, for the randomized alerting, had the RL agents running in the background, but both actions were ‘silent’ to the subject – and, in addition, there were random alerts played on some trials, explicitly unknown to the algorithms. Thus, the weights do not represent the same learned weights as in sub-figures A, B, C and D for contribution to the same task, though they do still give an indication of the relative importance of the different features to the activations of the first hidden layer’s neurons, based on the reaction time-based rewards.

**Figure 8:**
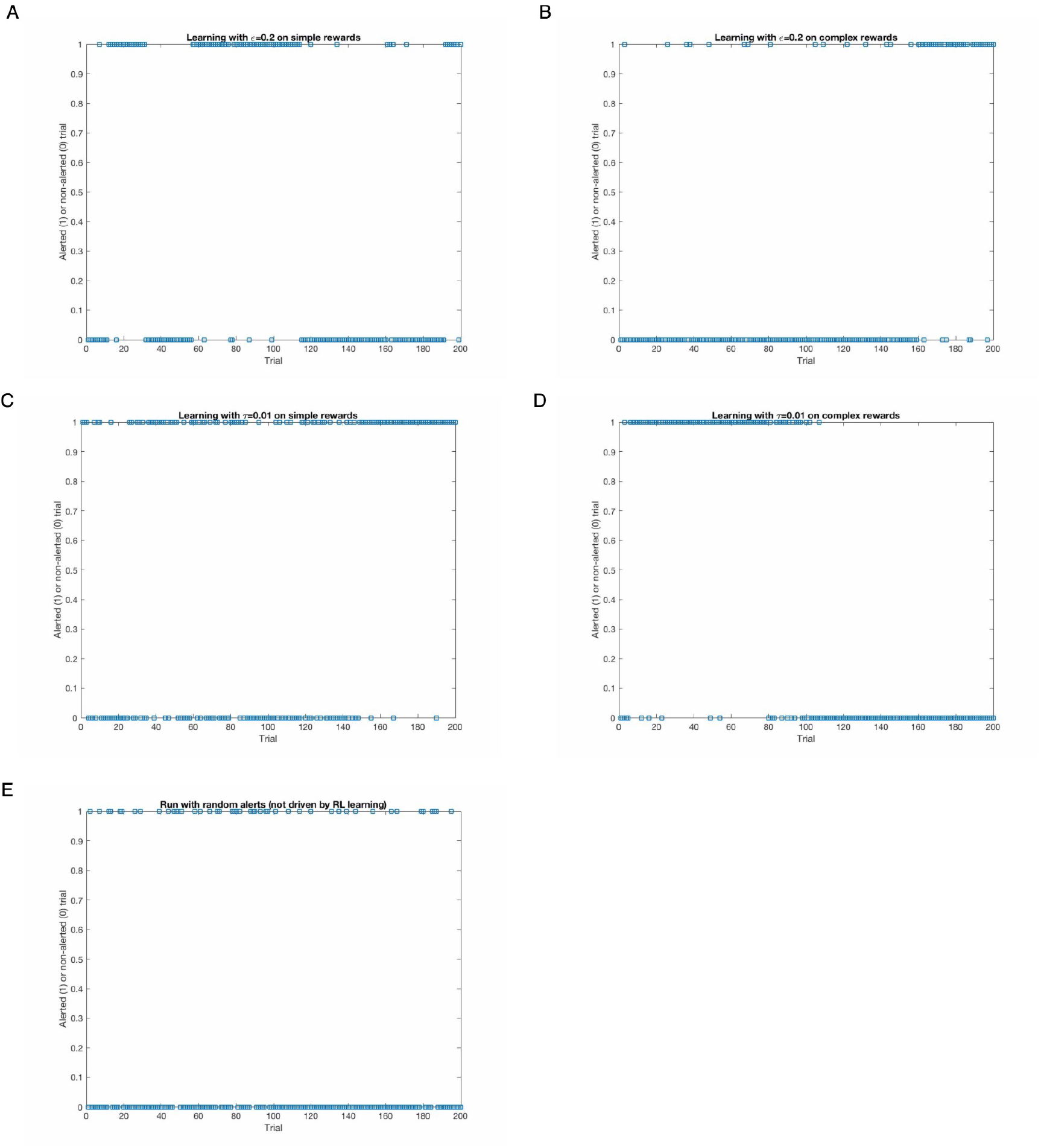
The figure shows the trials on which the RL system alerted (y value of 1) or did not alert (y value of 0) subject 1 for *ε*-simple (in A), for *ε*-complex (in B), for *τ*-simple (in C), for *τ*-complex (in D). E shows the randomized alerted vs non-alerted trials (RL ran in the background but did not control whether the subject was alerted or not).

**Figure 9:**
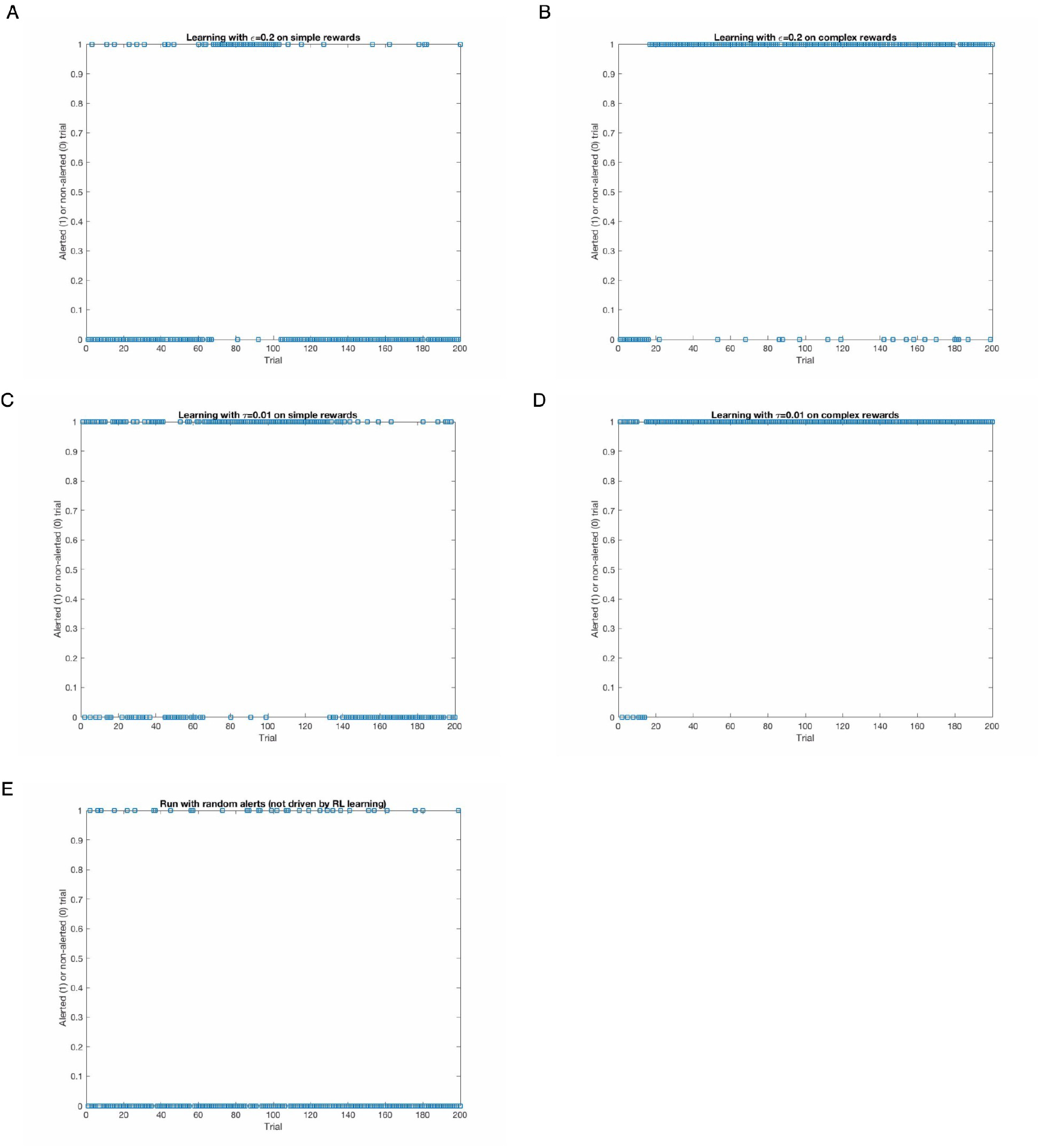
Analogously to Figure 8, this figure shows the trials on which the RL system alerted (y value of 1) or did not alert (y value of 0) subjet 2 for *ε*-simple (in A), for *ε*-complex (in B), for *τ*-simple (in C), for *τ*-complex (in D). E shows the randomized alerted vs non-alerted trials (RL ran in the background but did not control whether the subject was alerted or not).

**Figure 10:**
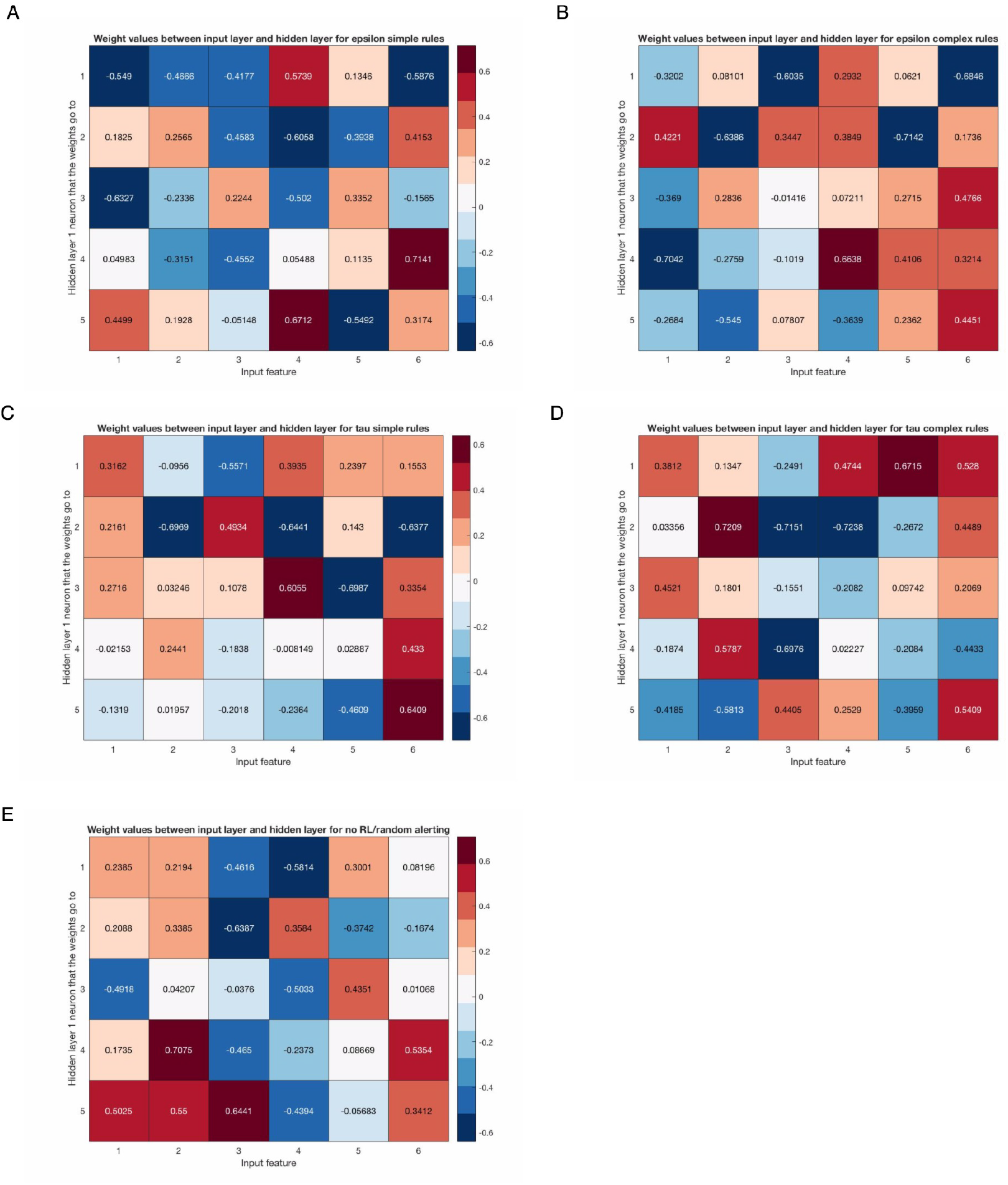
The figure shows the weights between the input layer and the first hidden layer of the DQN for subject 1, across all 5 conditions. We trained a new DQN for each condition, so the neurons in any of the layers do not directly correspond in any way, except that the input layer always has the same input features (the columns, described in the x-axis label). Feature 1 is the RT and the other 5 features are the linear relative band power values of the delta, theta, alpha, beta and gamma bands, respectively. There were 5 neurons in the first (and all other) hidden layers (hence the 5 rows). A shows these weights for *ε*-simple, B for *ε*-complex, C for *τ*-simple, D for *τ*-complex and E for the random alerts without RL. Note that for E, we show the weights for comparison. While the RL algorithm did run in the background, it did not alert – thus, it could not distinguish between action 1 (alerting) or action 2 (not alerting).

**Figure 11:**
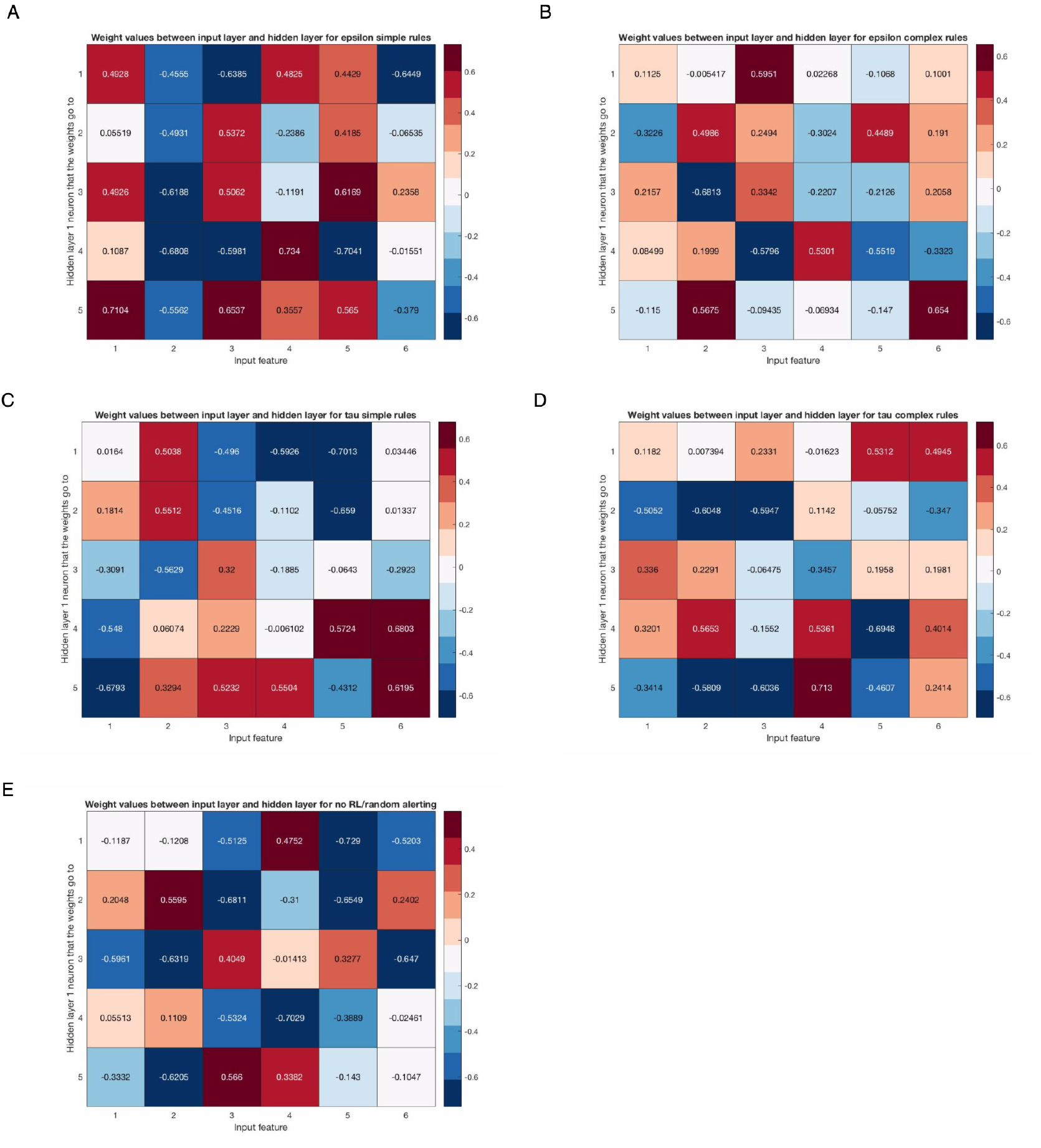
The figure shows, analogously to Figure 10, the weights from the input layer ot the first hidden layer neurons for subject 2. The sub-figures correspond to the conditions as in subject 1: A is for *ε*-simple, B for *ε*-complex, C for *τ*-simple, D for *τ*-complex and E for the randomized (non-RL-driven) alerts.

Next, we looked at the relationship between the reaction times and the rewards for the complex reward conditions. This had an almost perfect linear relationship for both subjects and this was almost perfectly identical for both action exploration strategies. We show a plot of the reaction times and the corresponding rewards for subject 1’s *ε-* complex run below:

Figure 12 suggests that a simpler reward rule where we reward with the negative of the reaction time may have achieved similar learning behavior. We discuss some of the issues around the reward function engineering in the discussion that follows.

**Figure 12:**
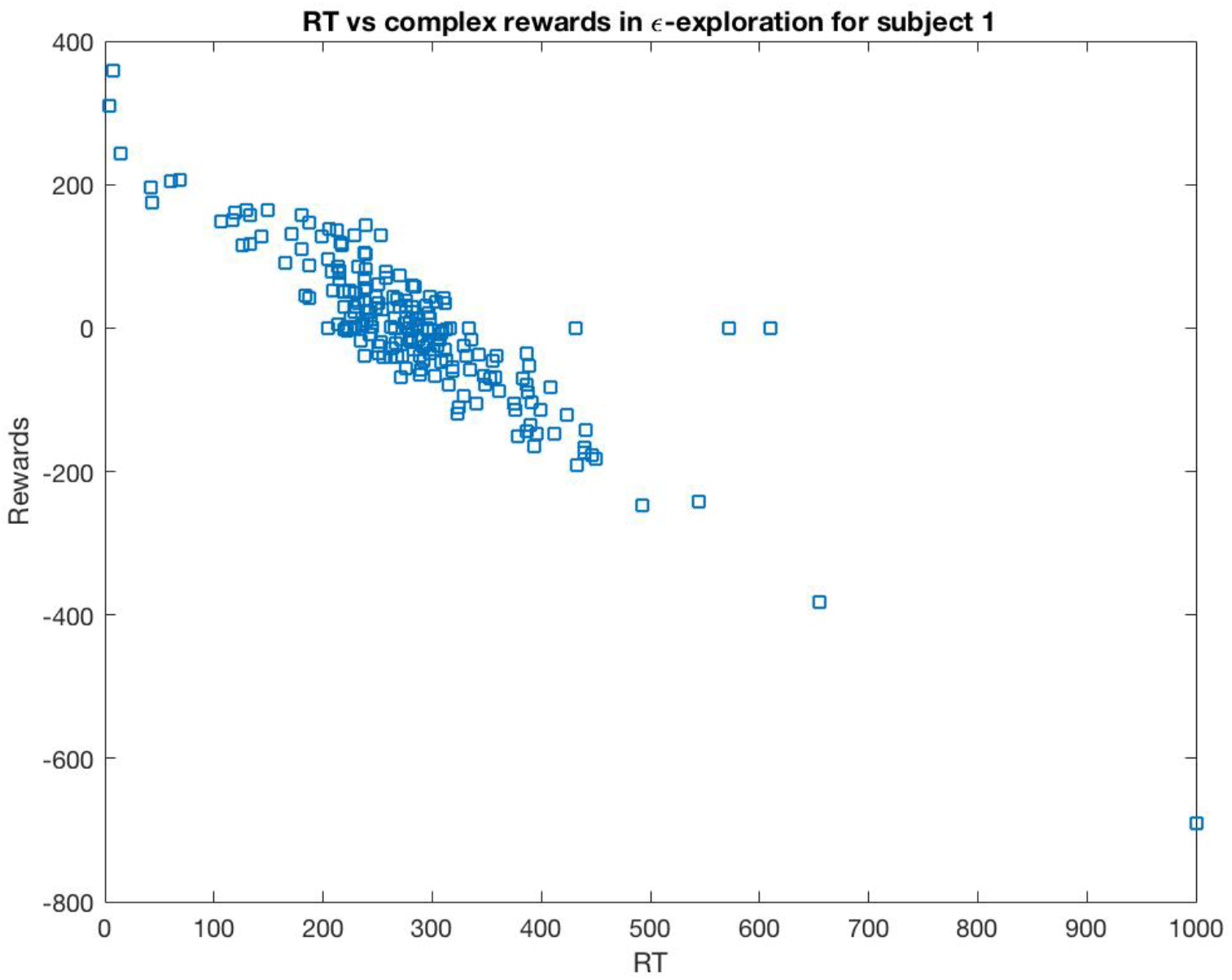
An illustrative plot of the more complex rewards (on the y-axis) vs the reaction times (on the x-axis) for *ε* exploration condition of subject 1. All other complex reward plots have the same form and strong linear relationship between the two. This relationship is not at all surprising of course, as the rewards are a function of how different the current reaction time was to the previous 10 trials’ mean RT. Since most trials will have RTs similar to the mean of the previous 10 trials, most differences (and therefore most rewards) will fall close to 0, which is what we see.

Finally, we looked for any linear relationships between the RTs and the relative EEG band powers, including interaction terms. All the p-values we report here are corrected for multiple comparisons using the Benjamini-Hochberg FDR procedure. For subject 1, we found no significant EEG band power predictors for the RT for *ε-* simple as well as the *ε*-complex and for the *τ*-complex conditions. For the *τ*-simple alerting, we find that all bands are slightly predictive of the RTs (p<0.05), as well as the interaction terms delta:theta power (p<0.05), delta:beta power (p<0.05), but most strongly alpha:gamma power (p<0.0005). There were other plausible significant effects before FDR-correction. Subject 2 had some significant p-values before FDR-correction but none after, for any of the conditions. We give some interpretation of these and the rest of the results next.

## 4. Discussion

We have presented here some preliminary results of using a powerful end-to-end neural network-driven approach to modeling both the brain state and the behavioral state, as well as when to play an auditory alert to the user of the system. Having shown that the system can learn in a simulation setting, with simulation parameters derived from empirical/experimental runs, we then run a (very) small scale pilot with 2 subjects. As this was a pilot study to test that the methods can work in this setting at all, our focus was not on deriving specific results related to the EEG and the reaction times. Our primary focus was to show that the system can learn non-random alerting behavior and, perhaps, alert better than a randomly-alerting system. This latter point was, in part, successfully shown empirically.

In our simulation results section, we mentioned and showed how Boltzmann-exploration of which action to choose leads to a more balanced alerting. Interestingly, for ‘good’ values of *τ* (i.e. around 0.01) the system agent learned to alert in a more similar proportion to *ε*-exploration, though still not as often. The latter, on the other hand, will pick the best action with probability 1-ε, which means that once the agent finds that the alerting state leads to a lower reaction time, it is likely to prefer and consistently use that action when choosing greedily. While for many situations this may be desirable, when alerting human users or operators, we do not want to alert constantly simply because alerting tends to improve the reaction time. This is why we implemented two reward rules, and showed results only with the approach which lead to a more balanced proportion of alerted trials. The two reward mechanisms tried were on the one hand where we reward either a 1 or a -1, or these multiplied by some factor (e.g. 0.1 and -0.1 or 0.3 and -0.3); and on the other hand where we reward as the difference between the mean of the moving window of the previous 10 states and the state we just ‘landed’ in. This has the effect of preferring to not alert too many times in a row, by virtue of the fact that if the system has just alerted many times in the window of states considered, then it is more likely that another alerted state will not have a lower reaction time and lead to a negative reward, or punishment (i.e. because of regression to the mean of the alerted states). A related reason for the ineffectiveness of too much alerting in a row is likely to be that the alerting system becomes habituated to the auditory alert. We have experimented with various types of implicit ‘regularizations’ or modifications to the reward function to encourage the system to not behave in one way for too long. One approach we explored and found useful is to add an ‘energy’ term to the agent, which gets decreased on alerting and increased on non-alerting to bias the system towards non-alerting. This makes understanding and analysis of the behavior of the system more complicated still, and the energy term needs to be adjusted depending on the expected value of the magnitude of the rewards for a given action.

Of course, running a continuous BCI NFB system is more complicated than the controlled case of the simulated distributions. Firstly, the brain and behavioral dynamics are highly non-stationary and would not follow the neat and non-changing distributions in the simulation (despite them being derived from empirical data). In general, while we expected alerted states to lead to lower reaction times, we found that this was not always the case. Part of the reason for this may be due to statistical reasons – i.e. too few alerted or non-alerted trials (e.g. subject 1 had too few alerted trials for the *ε*-complex condition and subject 2 had very few non-alerted trials for the *ε*-complex and *τ*-complex conditions). Unlike with the simulation results, finding that the agent was stuck in one behavior/action for many trials does not directly suggest that this is due to a convergence of the algorithm to the ‘best action’. Rather, we can imagine that being in one state for too long is likely to lead to diminishing returns (i.e. being alerted constantly for too long, or doing a repetitive and attention-demanding task for too long without the alerts) and the ‘optimal action’ is likely to switch. We observed this quite notably in the simple reward rules-driven alerting for both subjects. The complex rewards lead to a different behavior, where a single action was preferred more strongly over a longer continuous time (though switching to the other action eventually). Interestingly, the sliding window RT-difference reward rule (which is more complex than the simpler constant rewards) led to a simple and strong linear relationship between the ultimate reward and the reaction time – namely that the lower the reaction time, the higher the reward, with RTs around 0 having rewards around 0. This is not surprising though it is not at all immediately clear that simpler rewards of that form, where we reward the negative of the reaction time (i.e. small RTs are small punishments, large RTs are large punishments) or the inverse of the reaction time (small RTs lead to larger positive rewards and large RTs lead to smaller positive rewards), would lead to similar learning or behavior as the rule we employed. We found some plausible correlations of various degrees of certainty between the band powers (most notably the alpha band power and gamma band power linear interaction term) and the RT. We do not elaborate on those results as this work’s focus is on showing the ability to use this powerful deep neural network-driven RL approach to BCI and NFB.

An obvious weakness of the work we present here is that it is of a very small sample size (n=2). However, the point of the work is not so much to show generally-applicable neuroscientific results from this work directly, but rather to show that we can work with deep RL in a BCI/NFB setting, which we have shown. We have also shown the effects of changing some of the parameters on simulated and real data. While in general Boltzmann-exploration should learn better, as it exploits more information in the estimated Q values for the two actions, this is not necessarily observed. In part, this is because it is easy to pick default parameter values (e.g. *ε =* 0.1) for it which both quickly find the optimal state-action mapping while allowing for sufficient exploration, compared to Boltzmann-exploration, where sensible *τ* values depend on the specific Q-value magnitudes. These in turn depend on the reward function and therefore, in general, on the context to which the RL has been adapted to. It is very likely that Boltzmann action-randomization is going to be more beneficial when there are more actions that are weighted and thus there is more information between the relative estimated utility of the different actions compared to our binary case.

It is possible, and we found multiple times, trained agents to behave so as to maximize the rewards in ‘funny ways’. During development of the presented methods, we had a version of a reward function where a wrong trial, the current RT was set to a high value (e.g. 1000 or 2000 milliseconds). This had the effect of punishing the RL agent when the human subject made a mistake on the task (e.g. most commonly this was pressing the space key when the response should have been inhibited). Having this feature prompted agents with the complex reward rule to sometimes learn very peculiar and concerning alerting patterns under both *ε*-exploration and Boltzmann-exploration of the two actions, where they would alert most of the time and not alert only on rare occasions. This suggested to us that those agents were learning that alerted trials involve higher errors and thus higher punishments. Those agents would be silent (not alert) for a few trials approximately every ten trials (i.e. the width of the window). Since higher punishments mean that the last window of RTs compared to the current RT is likely to be a higher value, due to error trials being set to have effectively very high RTs. Thus, these agents seemed to prompt users to make more mistakes and then exploit the higher rewards resulting from the relatively much lower RTs on the few non-error trials after a sequence of constant alerting. This is obviously not optimal for the task at hand or for many tasks, but may lead to a higher overall cumulative reward for the agent, thus satisfying the problem it is solving, but not the problem we wanted it to solve.

An interesting application of this method might be to learn continuous actions that modify some set of parameters of a subject’s brain state. The actions could be, for example, the parameters of a transcranial alternating current stimulation (TACS) setup, such as the current, voltage, frequency, phase and the active electrodes in the stimulation setup. This is an area we hope to explore next with this methodology. However, we will conclude this work by highlight some of the dangers of relying on such methods for BCI, especially in settings where we affect the user or an external life or mission-critical system or affect their brain activity with an approach like TACS. Because it is very difficult to predict the details of how all the parameters interact with the reward rule in an RL system *a priori.,* we urge some awareness that the fact that a system learns to optimize for a certain reward rule does not mean that the system is behaving using desired action sequences in practice. Future BCI and NFB work employing complex RL-based or other complex statistical or ML-based methods should keep this mind.

